# Spatially resolved optical monitoring of spinal cord blood flow with a minimally invasive, multi-level epidural probe

**DOI:** 10.1101/2020.10.06.326900

**Authors:** David R. Busch, Wei Lin, Chia Chieh Goh, Feng Gao, Nicholas Larson, Joseph Wahl, Thomas V. Bilfinger, Arjun G. Yodh, Thomas F. Floyd

**Author notes:** No prior presentation. Corresponding author: Thomas F. Floyd, MD UTSW, Anesthesiology and Pain Management, Cardiothoracic Surgery, Radiology 5353 Harry Hines Blvd Dallas, TX 75390-9068 214/648-8235.

## Abstract

Spinal cord ischemia leads to iatrogenic injury in multiple surgical fields, and the ability to immediately identify onset and anatomic origin of ischemia is critical to its management. Current clinical monitoring, however, does not directly measure spinal cord blood flow, resulting in poor sensitivity/specificity, delayed alerts, and delayed intervention. We have developed an epidural device employing diffuse correlation spectroscopy (DCS) to monitor spinal cord ischemia continuously at multiple positions. We investigate the ability of this device to localize spinal cord ischemia in a porcine model and validate DCS versus Laser Doppler Flowmetry (LDF).

Specifically, we demonstrate continuous (>0.1Hz) spatially resolved (3 locations) monitoring of spinal cord blood flow in a purely ischemic model with an epidural DCS probe. Changes in blood flow measured by DCS and LDF were highly correlated (r=0.83). Spinal cord blood flow measured by DCS caudal to aortic occlusion decreased 62%, with a sensitivity of 0.87 and specificity of 0.91 for detection of a 25% decrease in flow. This technology may enable early identification and critically important localization of spinal cord ischemia.

---

Vascular surgery for management of aortic disease,^1–7^ and spine surgery for the management of congenital and acquired spine deformity^8, 9^ or spine trauma,^10^ not infrequently lead to iatrogenic spinal cord ischemia, which in turn may result in paralysis and paraparesis. To date, efforts to image the critical spinal cord blood supply preoperatively to guide surgical intervention have met with mixed success.^11–17^ The vasculature supplying the spinal cord is complex^18^ and exhibits large inter-subject variability^19, 20^ in both native vascular anatomy and as a result of prior aortic disease. These factors help to explain the frustration of efforts to prevent spinal cord ischemia.

The approaches most widely employed for intraoperative spinal cord ischemia monitoring are motor and somatosensory evoked potentials. Unfortunately, evoked potentials (EP) have high false positive and false negative rates,^21, 22^ and the changes in EP that drive alerts are often delayed by 10-20 minutes after ischemia onset.^23^ Additionally, EP have shown very limited ability to identify the anatomic origin of the ischemia. In clinical practice, electrophysiological monitoring is affected by anesthetics and hypothermia,^24^ is incapable of monitoring in the preoperative or immediate postoperative setting, and requires continuous presence of a technologist and/or neurologist. Thus, it is unclear whether current intraoperative neurophysiological monitoring in aortic^25^ or spine surgery^26^ influences outcome.

An optimized spinal cord monitor would offer the ability to immediately detect and regionally/axially resolve the level of origin of the ischemia, independent of anatomic variability in vascular supply. Such qualities could speed surgical response and focus it toward restoring spinal cord blood flow. To address these limitations, here we demonstrate a novel optical device to measure spinal cord blood flow. To measure spinal blood flow, the device employs diffuse correlation spectroscopy (DCS)^27, 28^ and utilizes a thin, highly flexible, epidural fiber optic probe with multiple flow sensors (FLOXsp). In a porcine model of pure ischemia (e.g., no associated trauma, compression, distraction), we test the ability of this device to rapidly detect and regionally discriminate changes in spinal cord blood flow using a single probe.

## A. Results

DCS data at three positions along the spinal cord were acquired at least once every 10 seconds throughout physiological manipulations and aortic inflations as described below for each of 5 pigs.

We utilized laser Doppler flowmetry (LDF) to measure blood flow at a site caudal to the lowest DCS sensor (~2 vertebral levels separation), placing a surface LDF probe directly on the spinal cord. During physiological perturbations designed to increase and decrease spinal cord blood flow (hypoxia + hypercarbia, or hypocarbia), the partial pressure of carbon dioxide in the blood (pCO_2_, measured by blood gas) ranged from 20.7 to 93.7 mmHg, inducing up to a ~20% decrease and ~200% increase from baseline in DCS-measured blood flow (Figure 1) while arterial blood oxygenation ranged from 100 to 38%. The changes in blood flow were highly correlated between the LDF and proximate DCS sensor (Figure 2).

**Figure 1:**
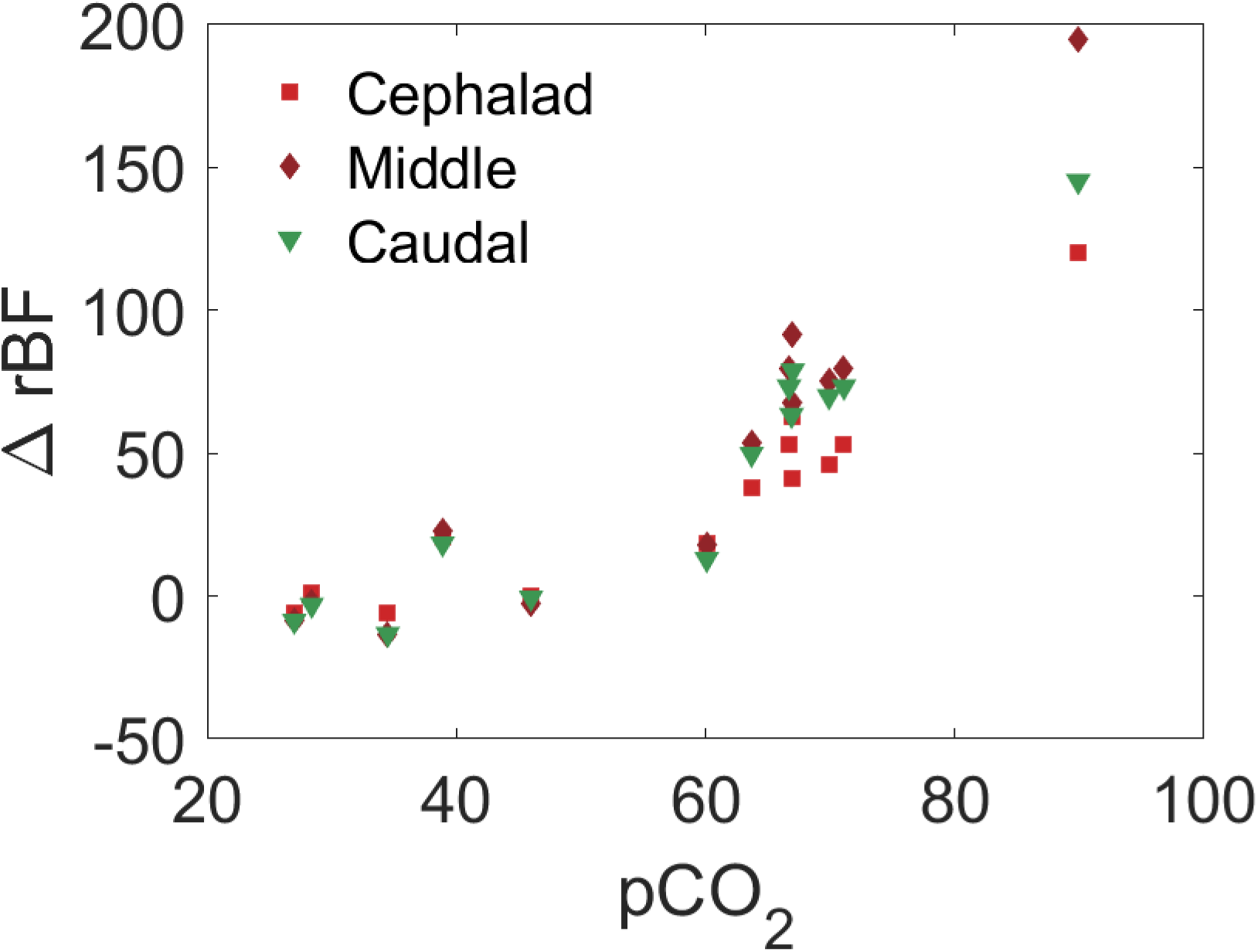
Example change in relative blood flow versus blood concentration of carbon dioxide measured by DCS at three sites along the spinal cord in a single pig.

**Figure 2:**
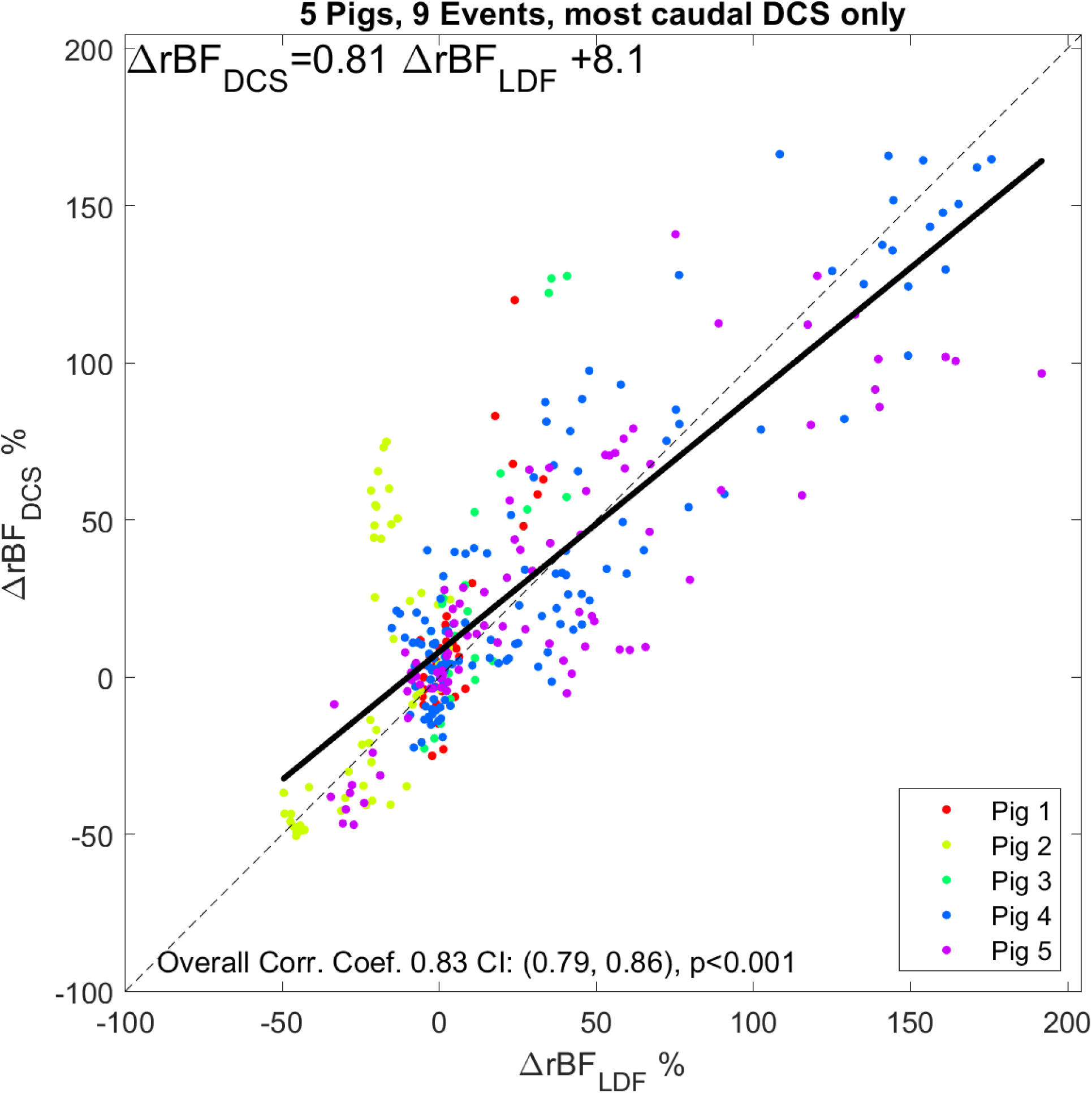
Comparison of changes in relative cerebral blood flow measured by the LDF and most caudal DCS sensors. A best linear fit line is shown (black) along with a 1:1 line (dashed black line). The overall correlation coefficient was 0.83 (p<0.001; 95% confidence interval, (0.79, 0.86)). Measurements made in 5 pigs during hypoxia-hypercarbia (pCO_2_ range 20.7 to 93.7 mmHg).

The balloon position in relationship to the FLOXsp probe regions and LDF probe was confirmed fluoroscopically prior to and during inflation, then monitored throughout the experiment. Aortic occlusion was confirmed by loss of femoral artery pressure. An example of a fully inflated balloon between the central and caudal DCS detectors is shown in Figure 3.

**Figure 3:**
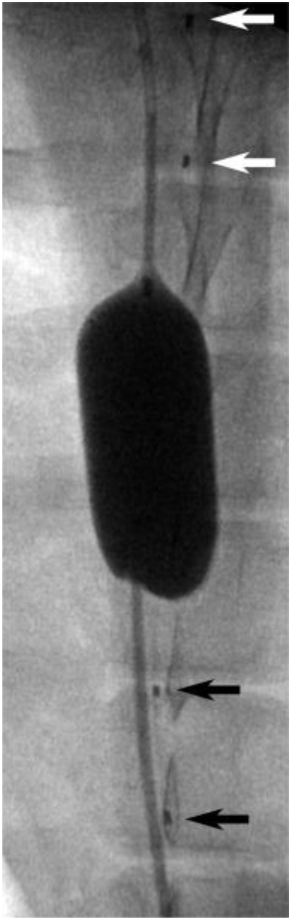
Anterior-posterior radiograph of an REOBA balloon inflated in the aorta between the caudal (black arrows) and central (white arrows) positions of a FLOXsp probe placed on the along the spinal cord in the epidural space. Fiducial markers noted by arrows are located between each light source and corresponding pair of detector positions (Figure 4).

The axial location of aortic occlusion, from the mid-thoracic to lumbar, led to regionally distinct changes in spinal cord blood flow. Detectors below the balloon occlusion level rapidly detected a decrease in blood flow, while detectors positioned above the inflated balloon detected quasi-stable or modestly increased spinal cord blood flow. Figure 4 depicts characteristic regional changes in blood flow sensed by the FLOXsp in response to balloon inflation. When the balloon is positioned cephalad to each sensor level (e.g., red shaded regions in Figure 4), flow decreases. In this example, we observed little or no change in flow in FLOXsp detector regions when the balloon was placed caudal to the sensor position (e.g., blue shaded regions in Figure 4). The more caudal the balloon was to the level of measurement, the less perturbation during balloon inflation was observed. Blood flow at FLOXsp detector regions caudal to the balloon inflation demonstrated reliable and substantial decreases in blood flow. Blood flow measured by LDF showed a similar pattern (Pearson’s correlation coefficient r=0.83), although the location of this sensor limited balloon positions cephalad to it. Increases in spinal cord blood flow immediately following balloon deflation were higher (up to +150%) in DCS measurements than in LDF (<50%) metrics. This response was not the focus of this study and the differences were not quantified, however, this observation suggests that blood flow in the deeper tissues (gray matter), able to be probed by DCS, may have different reperfusion dynamics than the relatively shallow tissues (white matter) probed by LDF.

**Figure 4:**
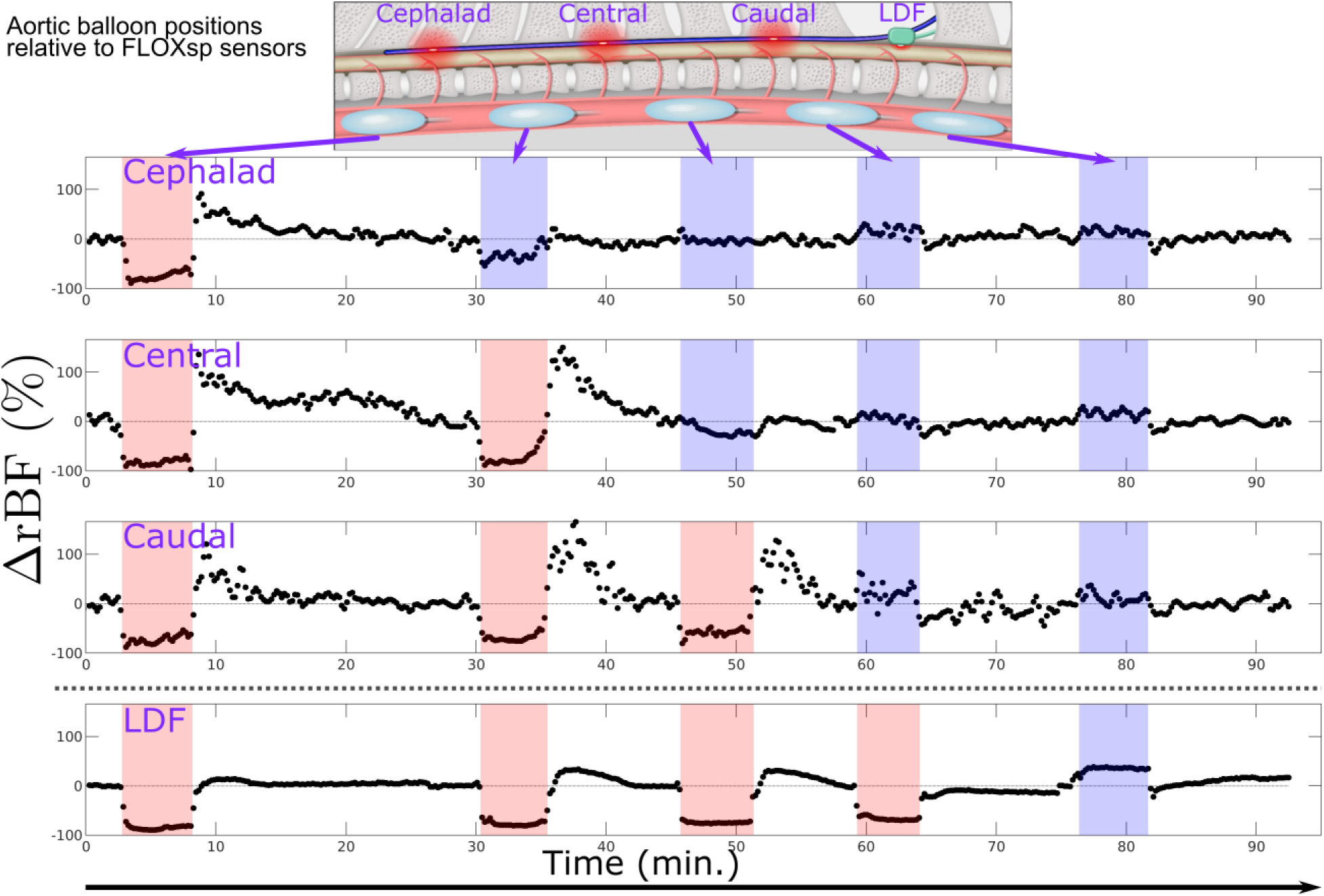
Time course of blood flow changes during serial aortic balloon inflation and occlusion, as the balloon is moved from cephalad to caudal locations (schematic, inset), measured at three different sites with DCS (cephalad, central, and caudal) and one site with LDF. Shaded regions indicate that the aortic balloon was inflated. Red (blue) shading indicates the balloon was cephalad (caudal) to the measurement site. In situations where the sensor position was only slightly *cephalad* to the balloon, we occasionally observed slight decreases (e.g., cephalad sensor near minute 30) and increases (e.g., caudal DCS near minute 60). This effect may be due to the precise local vascular anatomy. Anecdotally, we observed smaller increases in blood flow following balloon deflation with LDF, although this reactivity was not the focus of our study.

Specifically and more quantitatively, blood flow response in this group of 5 pigs measured cephalad to the balloon occlusion was slightly elevated 13 [−2 18] % (median, [IQR]), while mean blood flow caudal to the occlusion dropped by −62 [−75 −46] % from baseline (Figure 5a). We calculated a receiver operating characteristic curve for the ability of the FLOXsp probe to detect the location (caudal or cephalad) of aortic occlusion (Figure 5b). The FLOXsp device was found to have a sensitivity of 0.87 and specificity of 0.91 for detection of a 25% decrement in spinal cord blood flow below the level of aortic occlusion.

**Figure 5:**
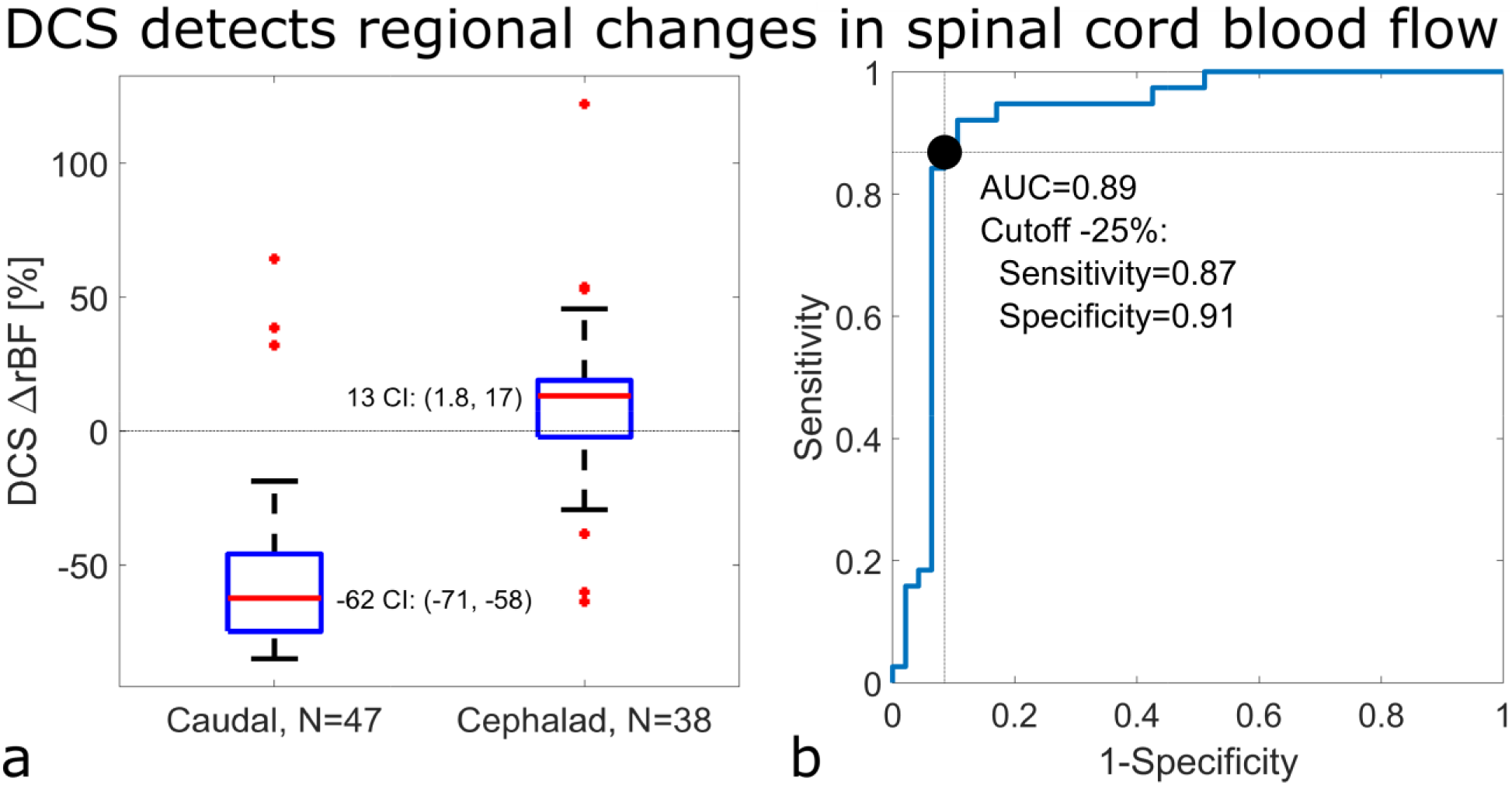
Blood flow changes in spinal cord regions cephalad and caudal to aortic occlusion at multiple positions and repetitions. No correction for multiple measurements in the same pig (N=5). (a) Blood flow below the site of aortic balloon inflation fell −62% (95% confidence interval, CI: −71, −58%) from baseline, while blood flow above the aortic occlusion increased 13 % (1.8, 17%) from baseline, rejecting the null hypothesis at a significance of 0.05 that the blood flow below the balloon site did not change (i.e., ΔrBF=0). The respective interquartile ranges for the caudal and cephalad measurements were [−75 −46] % and [−2 18] % respectively. (b) Receiver operating characteristic curve of the ability of DCS flow to predict the placement of the aortic balloon cephalad to the DCS sensors, including sensitivity (0.87) and specificity (0.91) calculated for a cutoff of ΔrBF=-25% to detect an aortic occlusion cephalad to the sensor location.

## B. Discussion

To test the ability of the device to detect and localize spinal cord ischemia we chose an approach that elicits ischemia without directly compressing the spinal cord. This was accomplished through use of an intra-aortic balloon which can be positioned and repositioned accurately using fluoroscopy between aortic occlusion (inflation) trials. Additionally, the probe is not disturbed during these periods of ischemia, minimizing potential motion artifacts due to the model. This approach also conveniently replicates aortic occlusion during open or endovascular thoraco-abdominal aortic repair.

The ability to regionally discern the origin of spinal cord ischemia is particularly important in repair of the thoracoabdominal aorta where the impact of the extent of resection or endovascular graft coverage remains unpredictable due to a high degree of variability in native vascular supply to the spinal cord, and/or to variability in effects of prior disease.^29, 30^ It is also of particular importance in the correction of spine deformity involving multiple levels where the origin of ischemia due to distraction may be uncertain. Currently, this enigma is compounded by the lack of reliability of evoked potential monitoring, including false negative and false positive alerts,^21^ as well as delays between inciting ischemia and alerts that can exceed 10 minutes.^23^ Our device has previously shown itself to rapidly (<10s) and reliably detect the onset of changes in blood flow.^31^ Moreover, evoked potential monitoring cannot localize ischemia, complicating any attempt to restore blood flow via intraoperative re-implantation, etc. In the present work, our device has demonstrated its ability to spatially localize the level of aortic occlusion, potentially offering an improved opportunity for focused intervention.

Our study had several limitations. We choose to use LDF as a comparator technique as it provides a continuous blood flow metric in the operating room. However, the LDF technique is sensitive to a small region (< 5 mm^3^) of superficial tissues (<2 mm from the surface), while the DCS technique as employed here is sensitive to a larger region (~ 5 cm^3^) up to ~10 mm into the tissue. Thus, comparison between the two methods is somewhat problematic as each samples a different tissue bed, likely partially explaining the discrepancy between techniques shown in Figure. Moreover, in the present experiment, the DCS and LDF probes were not co-located, as they both optically and mechanically interfere with one another. In addition, the LDF probe geometry created challenges in securing the probe in the laminotomy without injuring the spinal cord. LDF is known to be extremely sensitive to motion artifacts, especially if this changes fiber optic-to-tissue coupling. We utilized a temperature sensor built into the LDF probe head to identify and remove periods of poor tissue contact identified with lower temperature measurements.

Additionally, we excluded events in which the temporal derivative of the LDF signal had a discontinuity. DCS is also vulnerable to motion, however, the overlying bony spine effectively assists in holding the probe in place. DCS probe motion was not a challenge in this experiment; future long-term studies, especially if transport is involved, will need to address this issue.

## C. Outlook

In summary, the FLOXsp device is capable of immediate, continuous, reliable, and regionally specific detection of changes in spinal cord blood flow, offering earlier identification of spinal cord ischemia and improved opportunity for focused intervention. This device may prove of exceptional value to vascular and cardiovascular surgeons in planning for and conducting aortic interventions such as stenting, to spine surgeons in the management of congenital and acquired spine deformity, as well as in the management of spine trauma. Finally, while not yet tested in an intensive care environment, it is expected that this technology will also allow for continuous postsurgical monitoring, a critical period where monitoring is currently not possible and during which delayed ischemia resulting in injury is known to occur.

## D. Material and Methods

### D.1. Diffuse correlation spectroscopy (DCS)

DCS utilizes a diffusion model of light transport in tissue to provide rapid, quantitative, and non-invasive measurements of blood, as validated against several clinical techniques^27, 33, 34^ including in the spinal cord.^31^ DCS employs photon correlation techniques to derive a blood flow index proportional to microvascular blood flow.^27, 28, 35, 36^ Among other studies, DCS-monitoring of blood flow changes has been validated in humans against arterial spin-labeling and velocity mapping MRI.^33, 37–39^

The FLOXsp probe (Figure 6) is <0.13 cm in diameter, enabling placement into the epidural space through a small laminotomy or an epidural needle. The probe utilized in this study (FiberOptic Systems Inc, Simi Vally, CA) contained utilized 3 illumination fibers (100/110/125 μm LOH) separated by 10 cm, which were serially illuminated utilizing a 785 nm long-coherence laser (CrystaLaser, Reno, NV) through a fiber optic switch (PiezoJena, Jena, Germany). A single mode (6/125/165 Cu800) detector fiber is positioned on either side of each source with 2 cm of axial separation. Light was detected in parallel at these six locations using photon-counting APD detectors (Excelitas, Waltham, MA). Thus, the FLOXsp probe measures 3 distinct regions on the spinal cord, separated by 10 cm, with partially redundant measurements at each location (Figure 6).

**Figure 6:**
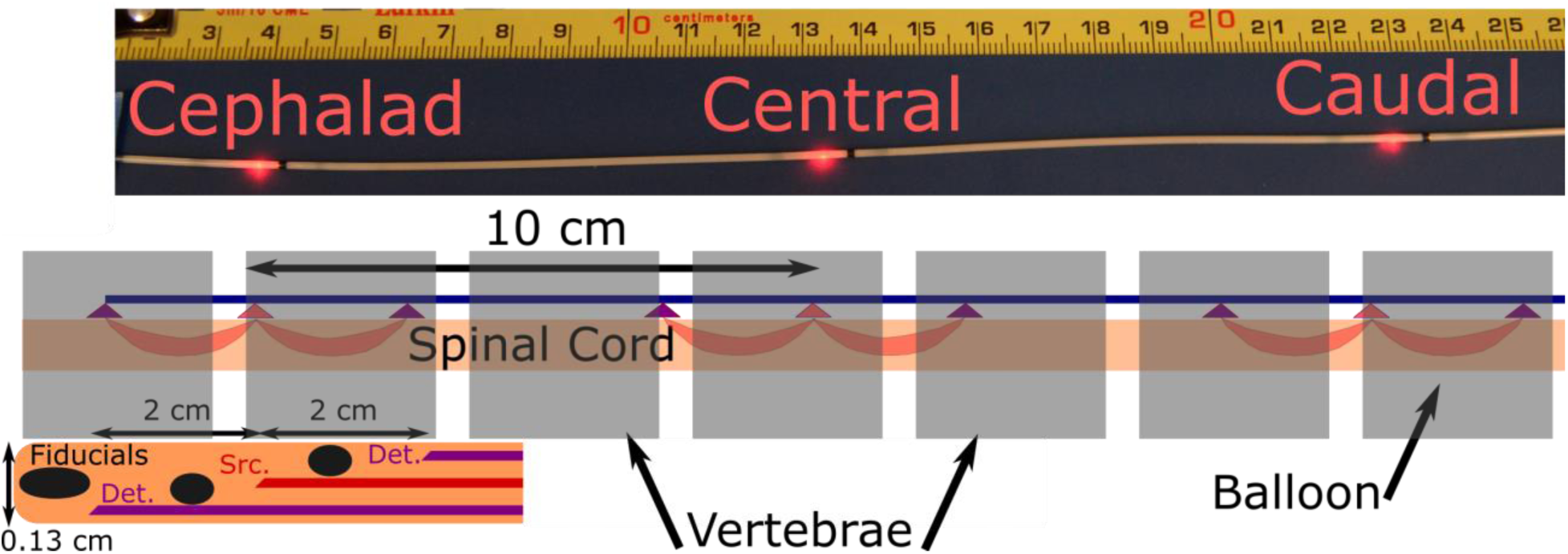
Photo of 3 position probe, all three source fibers illuminated. Black regions of probe are combined radiographic fiducials and blocks to prevent light propagation down the length of the probe. Schematic of probe in spine and expanded view of probe distal tip, showing fiducials at tip and between each source-detector pair, as well as the fiber positions. Each of the three detector-source-detector combinations along the probe is configured in a similar fashion. Not to scale.

Detector output was fed to a hardware correlator (Correlator.com) and the resulting temporal autocorrelation function transmitted to a data acquisition and instrument control computer with auxiliary data-acquisition and data-processing cards (PXIe-1082, PXIe-6612, PXIe-6361, PXIE-8135, National Instruments, Austin, TX). Here, data were fit in real-time to a solution of the correlation diffusion equation,^27, 28^ using an FPGA-based fitting routine described elsewhere;^40^ to arrive at a blood flow index (BFI), which is displayed in real time. Raw data was also stored for off-line processing. Data integration time and source intensity was adjusted for each animal; optical output power was limited to a maximum of 27mW, within the American National Standards Institute standard for skin illumination (no standard for spinal cord exists).^41^ Data acquisition of each ‘frame’, consisting of all three source positions, took <10s. In this study, we utilize changes in spinal cord blood flow, relative to a baseline region (ΔrBF(t)=(BFI(t)-BFI_baseline_)/BFI_baseline_), where the baseline region is prior to the experimental perturbation.

### D.2. Anesthesia and Monitoring

Five male and female domestic pigs weighing 47-65 kg were employed in these studies (Institutional Animal Care and Use Committee protocol 2018-102476-USDA). Anesthesia was induced with glycopyrrolate (0.2 mg/kg, IM), Ketamine (15-20 mg/kg, IM), Xylazine (1.1-2.0 mg/kg, IM) and Tiletamine HCL/Zolazepam (4-6 mg/kg, IM). Following anesthetic induction, a peripheral intravenous catheter was placed and animals were intubated and ventilation controlled. Anesthesia was maintained with isoflurane (1-3.0%, inhaled) and fentanyl (2-10 µg/kg/hr, IV). Central venous access was obtained using a 16 G single lumen catheter placed into the right internal jugular vein under ultrasound guidance and advanced into the right atrium to allow for reliable delivery of intravenous fluids and anesthetic agents. Adequacy of ventilation was verified via continuous end tidal CO_2_ and pulse oximetry, and intermittent arterial blood gas analysis. Rectal temperature was monitored and a Bair Hugger (3M Maplewood, MN) forced air warming blanket with an operating site orifice was also employed.

### D.3. Epidural Multi-level Probe Placement

The FLOXsp probe was placed into the epidural space through an L2/3 laminotomy, and then advanced into position with fluoroscopic guidance. The anatomic position (vertebral level) of each set of detectors was documented; the DCS sensors were positioned between the T7 and T9, T11 and T13, and T15 and L1 vertebrae, respectively (Figure 7).

**Figure 7:**
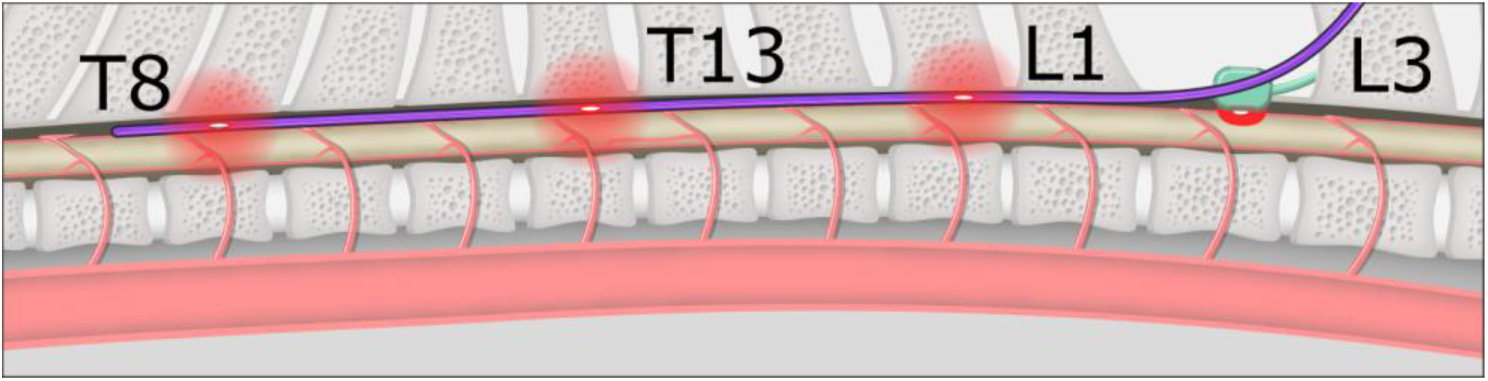
Schematic of pig spine with FLOXsp (blue) and LDF probes (green) in place. The REOBA balloon is shown in 5 positions, above/below each of the cephalad, central, and caudal DCS measurement sites (in this example, T9, T12, and T15), as well as the LDF site at the L2 laminotomy. Red clouds indicate the approximate zone of illumination for each sensor.

### D.4. Flow Validation and Regional Ischemia Detection

Laser Doppler Flowmetry (LDF) was employed to provide a continuous monitor of blood flow changes to compare to changes in spinal cord blood flow measured by DCS. While not approved for spinal cord monitoring, there is wide acceptance of LDF to measure superficial hemodynamics in other tissues, *e.g.*, brain.^42^ Using the same laminotomy, a laser Doppler flow (LDF) probe (CP1T-HP, Moor Instruments, Wilmington DE) was placed directly upon the spinal cord, caudal to the most proximal set of detectors on the FLOXsp probe (~2 spinal levels, Figure 7). The 8 mm diameter face of this probe was placed flat against the spinal cord, minimizing hemodynamic change due to the pressure of the probe on the spinal cord or mechanical injury of the cord. We compared changes in flow measured with the FLOXsp probe and LDF apparatus in two experiments:

1. <u>Physiological/Respiratory Manipulation of Flow:</u> Hypocarbia induced hypoperfusion and hypercarbia/hypoxia induced hyperemia. The effect of both hypocarbia and hypercarbia/hypoxemia upon neurovascular hemodynamics are well known. Hypocarbia was accomplished via step-wise increases in ventilation to reduce arterial pCO_2_. This was followed by stepwise decreases in ventilation to a final respiratory hold (maximum of 5 minutes or SpO_2_=50%) to achieve a peak level of pCO_2_ ~60 hhHg and hypoxemia (PaO_2_~45 mmHg). Pigs were then hyperventilated once again in a stepwise fashion to achieve hypocarbia (PaO_2_~25 mmHg). Arterial blood gases obtained during these experiments confirmed the extent of both hypo/hypercarbia.
2. <u>Multi-level Aortic Occlusion:</u> Here we correlated changes in regional blood flow measured by the FLOXsp probe and the LDF probe during aortic balloon occlusion. An ER-REBOA intra-aortic balloon catheter (Prytime Medical, Boerne, TX) was introduced via a 7 Fr sheath into the femoral artery. This balloon catheter also facilitates measurement of intra-aortic pressure, cephalad to the level of balloon inflation. The pigs were then placed in a prone position and an L-2/3 laminotomy was created. The FLOXsp probe was then placed into the epidural space through the laminotomy (schematic, Figure 7), advanced into position with fluoroscopic guidance, and the anatomic position (vertebral level) of each set of detectors documented. Following probe insertion, the REBOA balloon was advanced to the level of the proximal descending thoracic aorta and above the most distal detector set on the FLOXsp probe.

The balloon was then inflated to fully occlude the aorta at five locations: 1) cephalad to the distal FLOXsp detectors, 2) between the distal and central FLOXsp detectors, 3) between the central and proximal FLOXsp detectors, 4) between the proximal FLOXsp detectors and the LDF probe, and 5) below the LDF probe. At each location balloon inflation was verified fluoroscopically and with loss of femoral arterial pressure (Figure 3). Changes in flow were documented at each of the three FLOXsp detectors, as well as at the LDF probe. Balloon inflation was maintained at each position for 5 minutes.

#### Sample Size Determination

In determining sample size we assumed a non-normal distribution for the data and employed a one tailed Wilcoxon-Mann-Whitney test for two independent means. We assumed an ischemic region would experience a decrement of at least 25% in flow, and that the standard deviation in the measurement might approach 25%. Assuming a minimal effect size=1.0 for discriminating an injured from non-injured region and an α=0.05, sample sizes of 14 per group are be required to achieve power of 0.8-0.9. Given the ability to repeat the balloon inflation experiment several times in each pig, we estimated that a sample size of 8 or less would allow us to detect a difference in flow between regions. An interim analysis was conducted at 5 animals in an effort to reduce animal use and the data quality was deemed sufficient.

#### Statistical Approaches

DCS and LDF measurements of blood flow during hypoxia-hypercarbia were down-sampled to similar time bins, normalized to a baseline period prior to each manipulation of ventilation, pooled across pigs and time points, then correlated (corrcoef, Matlab 2019a, Mathworks). No correction was made for repeated measurements in the same pig or time-series measurements.

Blood flow in the spinal cord during aortic occlusions were averaged over 3 minutes after the balloon was fully inflated. Data from multiple pigs and serial inflations in the same pig were pooled. The median (prctile, Matlab 2019a) and 95% confidence intervals^43^ were calculated without statistical correction for non-independent data. A receiver operating characteristic (ROC) curve and area under the ROC curve (AUC) were calculated using average change from baseline of DCS-measured blood flow over 3 minutes following balloon inflation at each of the 3 DCS sensor locations, using radiographically-determined location of the aortic balloon and sensor as the ground truth (perfcurv, Matlab 2019a).

## E. Acknowledgements

The authors thank the staff of the UTSW Animal Resource Center for assistance, Dr. Owoicho Adogwa for clinical insight, and Dr. Ashwin Parthasarathy and Kenneth Abramson for technical discussions.

This study was funded by the United States National Institutes of Health (U01-NS095761, secondary support from R01-NS060653).

Several authors have patents issued US20140343384 (AGY, TFF), US8,082,015 (AGY), US6,076,010 (AGY), US10,342,488 (AGY, DRB) or pending PCT/US2015/017286 (AGY, DRB) relative to DCS technologies. None of these patents are licensed or generate royalties.

Dr. Floyd is the CEO and President of, and Drs. Floyd, Lin, and Busch hold equity interest in, NFOSYS, Inc. This company may license the technology used in this study. There is currently no license and no sales.

## F. Author Contributions

Study conception and design: DRB, AGY, TVB, TFF; method and technological development: DRB, CCG, WL, AGY, TFF; study conduct: DRB, FG, NL, JW, TFF; data analysis and interpretation: DRB, CCG, WL, TVB, AGY, TFF; manuscript approval: all authors.

